# Mitochondria Regulate the Cell Fate Decisions of Megakaryocyte-Erythroid Progenitors

**DOI:** 10.1101/2024.11.17.623989

**Authors:** Eunkyu Sung, Shohei Murakami, Masanobu Morita, Tomoaki Ida, Takaaki Akaike, Hozumi Motohashi

## Abstract

Recent studies highlight the critical role of mitochondria in hematopoiesis, especially in stem cell function and erythroid maturation. To explore mitochondrial contributions to cell lineage commitment of hematopoietic progenitors, we utilized *Cars2* mutant mice, an ideal model for this purpose. CARS2, a mitochondrial isoform of cysteinyl-tRNA synthetase, has cysteine persulfide synthase (CPERS) activity. Our new mouse model, with reduced CPERS activity, showed that the *Cars2* mutation led to mitochondrial inhibition and anemia by suppressing erythroid commitment in megakaryocyte-erythroid progenitors (MEPs). This suppression was reproduced using mitochondrial electron transport chain inhibitors. We identified two distinct MEP populations based on the mitochondrial content: mitochondria-rich MEPs favored erythroid differentiation, while the mitochondria-poor MEPs favored megakaryocyte differentiation. These findings reveal critical contributions of mitochondria to the MEP lineage selection, acting as a “mitochondrial navigation” for lineage commitment.

## Introduction

Lineage specification in hematopoiesis has been extensively studied, particularly in the context of lineage-specific transcriptional regulation (Sive & Göttgens, 2014; Laslo et al., 2006; Kato et al., 2018; Murakami et al., 2014; Yu et al., 2017; Johnsonet al., 2022). Hematopoietic stem cells (HSCs) give rise to hematopoietic progenitor cells (HPCs), which are crucial for daily hematopoiesis as they generate functionally mature cells under steady-state conditions (Sun et al., 2014). These HPCs include three major populations: common myeloid progenitors (CMPs), megakaryocyte-erythroid progenitors (MEPs) and granulo-monocyte progenitors (GMPs) (Akashi et al., 2000; Yamamoto et al., 2013). Key transcriptional regulators have been identified for MEPs and GMPs as GATA1 and PU.1, respectively (Hoppe et al., 2016; Arinobu et al., 2007). Within MEPs, KLF1 and FLI1 are crucial transcription factors that drive differentiation into erythroblasts and megakaryocytes, respectively (Starck et al., 2003; Doré & Crispino, 2011). However, limited attention has been devoted to investigating the specific roles of intracellular organelles in lineage determination.

Mitochondria, often referred to as the cell’s “powerhouse,” are crucial for generating ATP, which is vital for maintaining cellular homeostasis and survival. Beyond energy production, mitochondria are essential for the biosynthesis of key macromolecules like lipids, heme, and iron-sulfur clusters, highlighting their function as a signaling organelle (Chandel, 2015). These functions place mitochondria at the center of crucial biological processes, including cell proliferation, differentiation, and stress adaptation. Traditionally, cellular actions are thought to be directed by the nucleus, with mitochondrial changes seen as secondary. However, recent studies have demonstrated that changes in mitochondrial metabolism can significantly influence the fate and function of HSCs in a manner similar to how transcriptional networks regulate the stem cell destiny. For example, HSCs with high content of mitochondria display long-term repopulation activity (de Almeida et al., 2017), and the mitochondrial electron transport chain (ETC) function and fatty acid oxidation play key roles in regulating HSC fate *in vivo* (Ansó et al., 2017; Ito et al., 2012; Yu et al., 2013). In addition, mitochondria are essential for the maintenance and functionality of mature hematopoietic cells. Specifically, mitochondrial function is vital for erythroid maturation, particularly in processes such as heme synthesis, iron metabolism and iron-sulfur cluster biogenesis (Fontenay et al., 2006; Ducamp and Fleming, 2019). Genetic studies in mice have further demonstrated that mitochondrial protein synthesis is required for the development and maturation of erythroid cells (Liu et al., 2017; Gotoh et al., 2020). Human cases with mutations in mitochondrial tyrosyl-tRNA synthetase (YARS2), mitochondrial leucyl-tRNA synthetase (LARS2) and mitochondrial isoleucyl-tRNA synthetase (IARS2) have been documented (Riley et al., 2010; Riley et al., 2015; Gong et al., 2023). These cases presented with sideroblastic anemia, an indicator of impaired heme synthesis, suggesting that mitochondrial translation is essential for proper heme synthesis. Despite the recognized importance of mitochondria in determining HSC fate, function, and post-lineage development, their role in the lineage selection of HPCs remains to be fully elucidated.

Supersulfides, diverse molecular species that are characterized by sulfur catenation, have recently been identified as conserved biomolecules across species (Murakami et al., 2023; Barayeu et al., 2023; Akaike et al., 2024). These molecules play key roles in various biological processes, including redox balance control, inflammation regulation, energy metabolism and hypoxic response (Ida et al., 2014; Zhang et al., 2019; Takeda et al. 2023; Sekine et al., 2024). Cysteine persulfide, a primary supersulfide, is generated in *de novo* synthesis pathways, with multiple enzymes exhibiting cysteine persulfide synthesizing activity. We recently found that mitochondrial cysteinyl-tRNA synthetase (CARS2) has a moonlighting function as a cysteine persulfide synthase (CPERS) (Akaike et al., 2017). The CPERS activity of CARS2 plays a critical role in the maintenance of mitochondrial energy metabolism, including ETC function, polarization of mitochondrial membrane potential (MMP), oxygen consumption, and ATP production (Akaike et al., 2017; Alam et al., 2023).

To elucidate the role of mitochondrial function supported by the CPERS activity of CARS2, we generated a new inducible *Cars2* mutant mouse model and analyzed their phenotypes and mitochondrial activity. Following the induction of *Cars2* locus recombination, the mice exhibited anemia-like phenotypes. Our findings revealed that the lineage choice of MEPs toward erythroid cells was blocked in *Cars2* mutant mice whose mitochondrial activity was suppressed as we expected. Consistent with these results, inhibitors of mitochondrial ETC function impeded the lineage choice of wild-type MEPs toward erythroid cells. Additionally, we discovered that the mitochondrial content determines the lineage choice of MEPs, indicating that mitochondria play a crucial role in the cell fate determination of MEPs.

## Results

### *Cars2* IKO mice exhibit anemia-like phenotype associated with erythroblast reduction

To investigate an *in vivo* contribution of CARS2-mediated CysSSH synthesis, we used a mouse line harboring *Cars2*^AINK^ mutation, which attenuates the CPERS activity but retains the cysteinyl-tRNA synthesizing activity (Akaike et al., 2017). Because *Cars2*^AINK/AINK^ mice were embryonic lethal (Matsunaga et al., 2023), we established an inducible system by combining *Cars2*^AINK^ allele and a floxed allele of *Cars2* (*Cars2*^F^) (Figure S1A and S1B) in the presence of tamoxifen-inducible Cre (CreER^T2^). To label cells that were subjected to recombination or to label donor cells in transplantation experiments, LSL-tdTomato allele was also combined. *Cars2*^AINK/F^:*Rosa*-CreER^T2^ (:*Rosa*-LSL-tdTomato) (*Cars2* IKO) and *Cars2*^+/F^:*Rosa*-CreER^T2^ (:*Rosa*-LSL-tdTomato) (*Cars2* IC) were compared in this study. Before and after 3-day treatment of tamoxifen, body weight and peripheral blood were examined on day 7 and day 14 (Figure S1C). While body weight was unchanged in both mice (Figure S1D), red blood cells (RBCs), hemoglobin (HGB) and hematocrit (HCT) were significantly decreased in *Cars2* IKO mice (Figures 1A and S1E). In contrast, the *Cars2*^AINK^ mutation increased platelets (PLTs) with a marginal effect on white blood cells (WBCs) (Figures 1A and S1E). Recombination efficiency was verified to be over 80% in both mice (Figures S1F).

**Figure 1.**
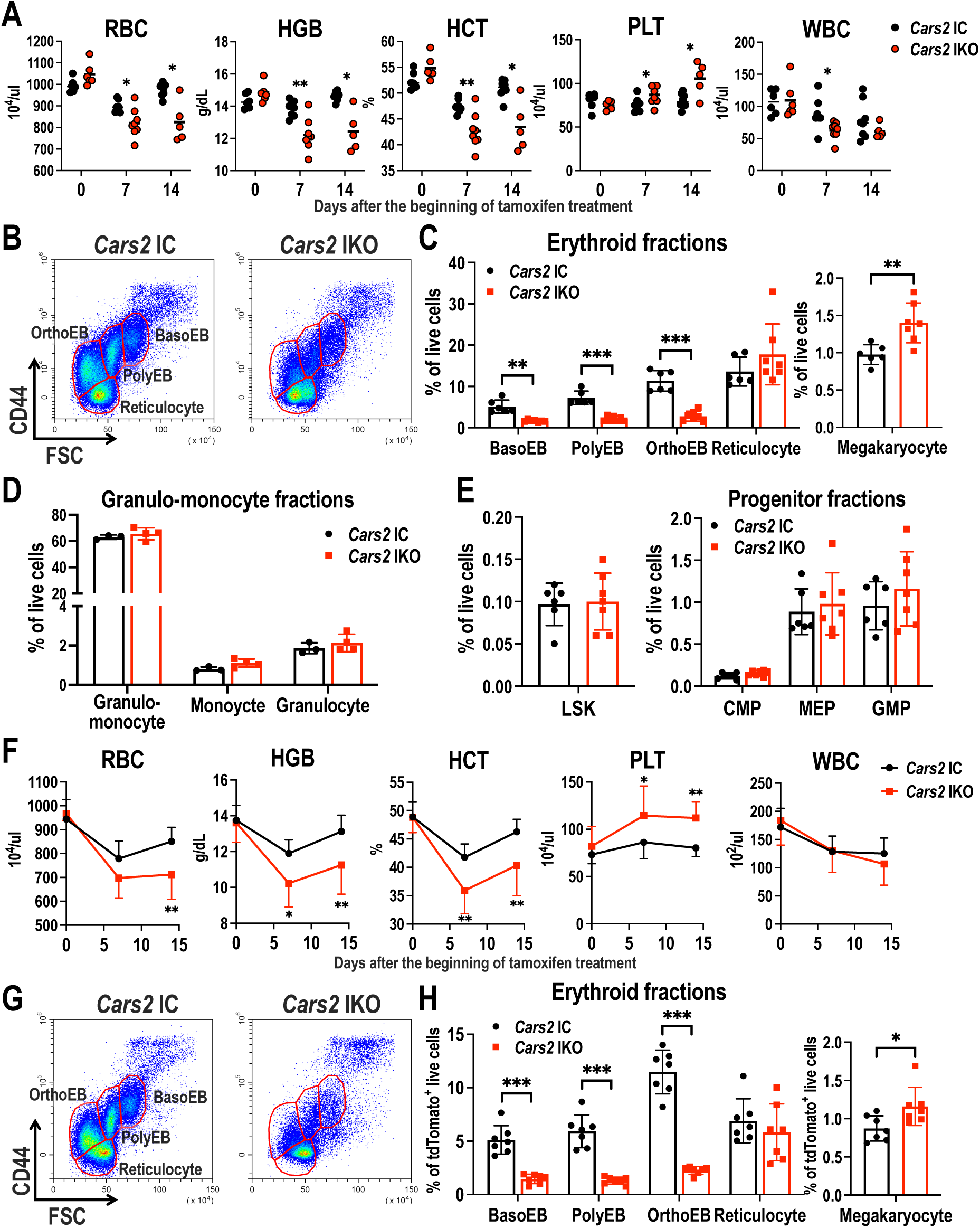
*Cars2* IKO mice exhibit anemia-like phenotype associated with erythroblast reduction. (A) Red blood cell (RBC), hemoglobin (HGB), hematocrit (HCT), platelet (PLT) and white blood cell (WBC) in peripheral blood at the indicated days. (B) Representative plots of CD44 versus FSC in Ter119^+^ BM cells. (C-E) BM profiles. Percentages of erythroblasts/megakaryocytes (C), myeloid cells (D), LSK cells and progenitor fractions (E). (F-H) Bone marrow reconstitution in recipient mice after non-competitive transplantation of *Cars2* IC and *Cars2* IKO BM cells (*n* = 7-10). Blood count parameters in peripheral blood (F), representative plots of CD44 versus FSC in Ter119^+^ BM cells (G) and percentages of erythroblasts/megakaryocytes (H). Values are presented as the means ± SD. Two-sided Welch’s *t*-test (A, C-E and H) and two-way ANOVA (F) were conducted to evaluate statistical significance. *P < 0.05; **P < 0.01; ***P < 0.001.

We next examined BM cells. In *Cars2* IKO BM, an erythroid fraction (CD71^+^Ter119^+^ (R2)) was reduced compared to *Cars2* IC BM (Figures S1G and S1H). Detailed analysis of erythroid differentiation stages (Figure S1I) revealed that the *Cars2*^AINK^ mutation reduced basophilic erythroblasts (BasoEBs), polychromatic erythroblasts (PolyEBs), and orthochromatic erythroblasts (OrthoEBs) (Figures 1B and 1C). In contrast, megakaryocyte fraction (CD41^+^) was increased in *Cars2* IKO BM (Figure 1C). No significant difference was observed in myeloid or lymphoid fractions (Figure 1D and S1J). Hematopoietic stem and progenitor cell (HSPC) fractions, Lineage^−^Sca1^+^cKit^+^ (LSK) cells, CMPs (Lineage^−^Sca1^−^cKit^+^CD34^+^FcγR^−^), GMPs (Lineage^−^Sca1^−^cKit^+^CD34^+^FcγR^+^) and MEPs (Lineage^−^Sca1^−^cKit^+^CD34^−^FcγR^−^), were also hardly affected (Figure 1E and S1K). Non-competitive bone marrow transplantation (BMT) (Figures S2A and S2B) showed that the peripheral blood and BM profiles of the recipient mice reproduced those of the donor mice (Figures 1F-1H and S2C-S2E), indicating that the *Cars2*^AINK^ mutation alters the hematopoiesis in a cell-autonomous manner, specifically the production of erythroblasts and megakaryocytes.

Since cell death is one of the causes for the reduction in cell fractions, we examined apoptosis of erythroblasts and megakaryocytes in BM cells and found that both pro-apoptotic and apoptotic cell fractions were rather decreased in *Cars2* IKO compared to *Cars2* IC mice (Figure S2F and S2G), suggesting that the reduction in erythroblasts due to *Cars2*^AINK^ mutation is less likely to result from cell death.

### Differentiation from MEPs into erythroblasts is impaired in *Cars2* IKO

Because erythroblasts and megakaryocytes were reduced and increased, respectively, in *Cars2* IKO mice without reduction in their progenitors, MEPs, (see Figure 1E), and because the reduction in erythroblasts is less likely to due to cell death (see Figures S2F and S2G), we focused on the binary differentiation capacity of MEPs in *Cars2* IKO mice.

To assess *in vitro* differentiation capacity of MEPs, we first conducted a colony-forming cell assay using Lineage^−^Sca1^−^cKit^+^ (LK) cells containing MEPs and GMPs. BFU-E formation from *Cars2* IKO cells was severely impaired, whereas CFU-M/G/GM formation was less affected (Figure 2A), supporting the idea that MEPs are selectively dysfunctional due to the *Cars2*^AINK^ mutation. Importantly, *Cars2* IKO MEPs were as proliferative as *Cars2* IC cells (Figure 2B), ruling out the possibility that the reduction in *Cars2* IKO BFU-E was due to a proliferation defect of *Cars2* IKO MEPs.

**Figure 2.**
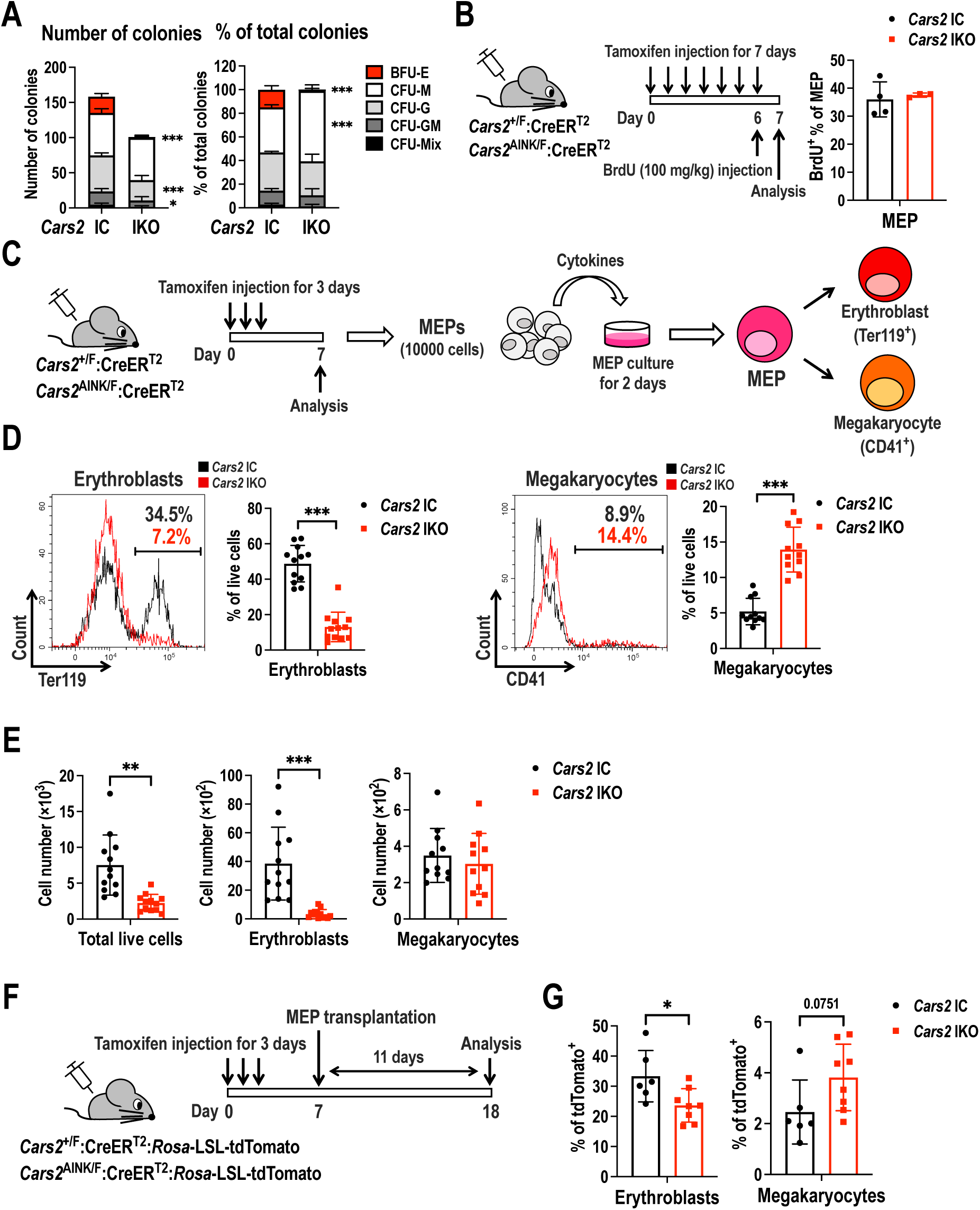
Erythroblast production is compromised in the MEPs of *Cars2* IKO mice. (A) Colony counts (left) and ratios (right) of each colony type in colony-forming cell assay (*n* = 3-4). BFU-E, burst-forming unit-erythroid; CFU-M, colony-forming unit-monocyte; CFU-G, colony-forming unit-granulocyte; CFU-GM, mixed colonies including CFU-G and CFU-M; CFU-Mix, mixed colonies including CFU-G, CFU-M and BFU-E. (B) Schematic illustration for BrdU incorporation experiment (left). Percentage of BrdU-incorporated MEPs (right). (C) Schematic illustration of *in vitro* MEP differentiation experiment. MEPs were cultured with cytokines (EPO, TPO, SCF, IL-3, IL-6) for 2 days and analyzed for erythroblasts (Ter119^+^) and megakaryocytes (CD41^+^). (D, E) Erythroblasts and megakaryocytes generated in the differentiation culture of MEPs. Representative histograms and percentages of erythroblasts and megakaryocytes (D) and counts of total live cells, erythroblasts and megakaryocytes (E). (F) Schematic illustration of MEP transplantation. MEPs were sorted after tamoxifen injection and transplanted into non-irradiated wild-type mice. Bone marrow from the recipient mice, which are labeled with tdTomato fluorescence, was analyzed on day 11 after transplantation. (G) Percentages of tdTomato^+^ cells in erythroblasts (Ter119^+^) and megakaryocytes (CD41^+^). Values are presented as the means ± SD. Two-way ANOVA (A) and two-sided Welch’s *t*-test (B, D, E and G) were conducted to evaluate statistical significance. *P < 0.05; **P < 0.01; ***P < 0.001.

We next assessed the differentiation capacity of MEPs using an *in vitro* culture assay, in which MEPs are differentiated into both erythroblasts and megakaryocytes. MEPs were cultured in liquid medium with cytokines for 2 days and analyzed for erythroblast and megakaryocyte differentiation using the surface markers, Ter119^+^ and CD41^+^, respectively (Figure 2C). The erythroblast and megakaryocyte ratios in *Cars2* IKO MEP culture were lower and higher, respectively, than those in *Cars2* IC MEP culture (Figure 2D). Notably, total cell and erythroblast counts were decreased in *Cars2* IKO MEP culture, whereas megakaryocyte counts were almost comparable between the two genotypes (Figure 2E), suggesting that *Cars2* IKO MEPs can produce megakaryocytes but almost lose the ability to produce erythroblasts.

To further evaluate the differentiation capacity of MEPs *in vivo*, we conducted a MEP transplantation, which we newly established to evaluate the MEP function (Figure 2F). MEPs were transplanted into wild-type recipients without irradiation. After 11 days, donor-derived tdTomato-positive cells were analyzed. Consistent with the *in vitro* MEP culture assay, fewer erythroblasts and more megakaryocytes were generated from *Cars2* IKO MEPs than *Cars2* IC MEPs, although the latter did not reach statistical significance (Figure 2G). Thus, the *Cars2*^AINK^ mutation inhibits erythroid commitment of MEPs without losing the potential to proliferate and differentiate to megakaryocytes.

### Mitochondrial dysfunction in *Cars2* IKO MEPs

Since the *Cars2*^AINK^ mutation is critical for mitochondrial function (Akaike et al., 2017; Matsunaga et al., 2023), we expected mitochondrial function to be impaired in *Cars2* IKO cells. Although mitochondrial mass was not altered by the *Cars2*^AINK^ mutation (Figure 3A), MMP and ATP/ADP ratio were reduced in *Cars2* IKO MEPs (Figures 3B-D), suggesting that the mitochondrial electron transport chain (ETC) is affected. Indeed, subunits of ETC complexes were reduced in *Cars2* IKO MEPs (Figure 3E). To investigate a functional impact of the observed changes in the ETC, we measured reactive oxygen species (ROS), as ROS have been implicated in mitochondrial dysfunction. Mitochondrial ROS were remarkably higher in *Cars2* IKO MEPs than in *Cars2* IC MEPs, whereas the intracellular ROS levels were comparable (Figures 3F and 3G). These results suggest that the mitochondrial function is substantially impaired in *Cars2* IKO MEPs.

**Figure 3.**
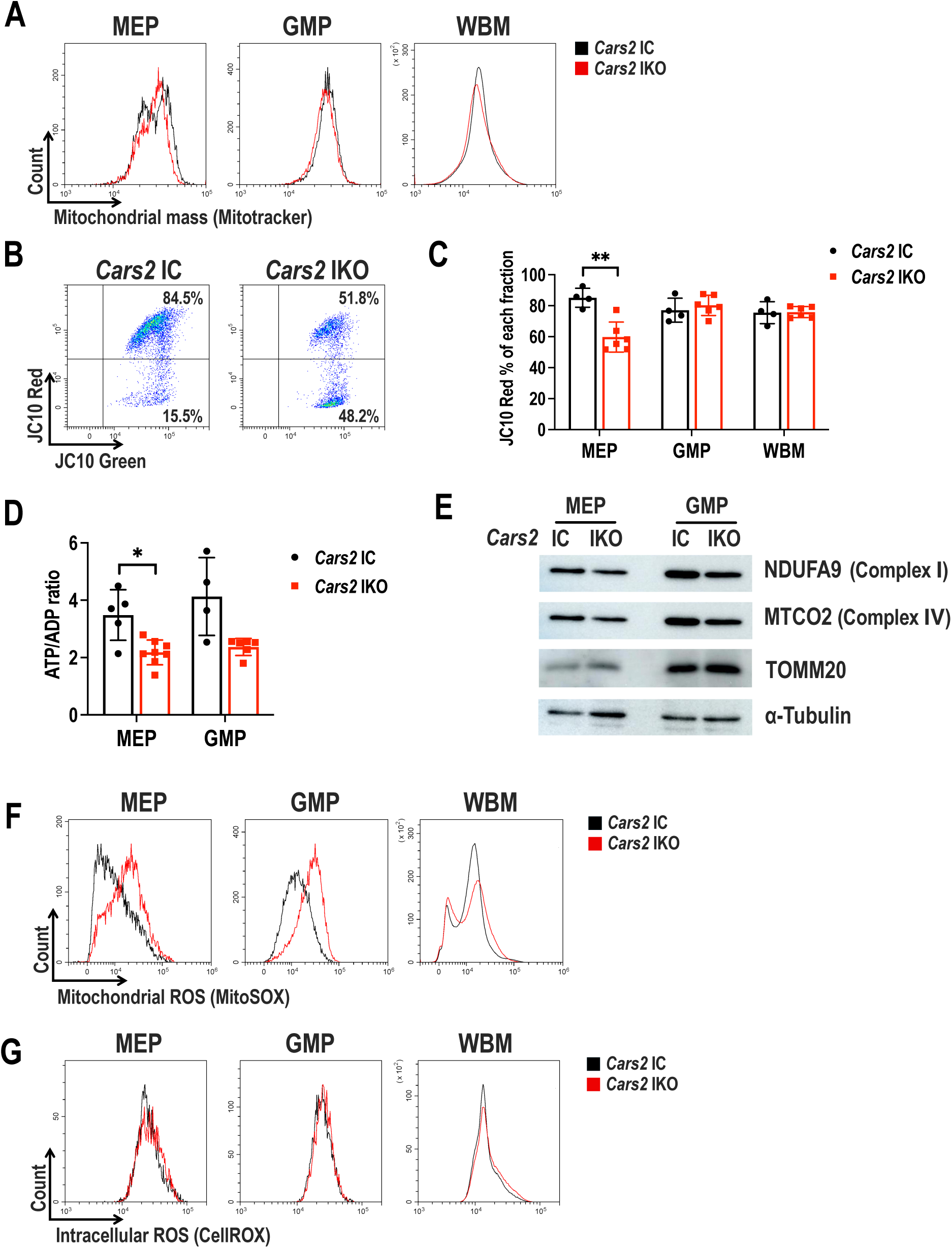
Mitochondrial dysfunction in *Cars2* IKO MEPs. (A) Representative histograms showing mitochondrial mass in MEPs, GMPs, and whole bone marrow (WBM) cells. (B, C) Mitochondrial membrane potential (MMP) in MEPs, GMPs and WBM cells. Representative plots of JC10 staining in MEPs (B). Ratios of JC10-Red-positive cells (C). (D) ATP/ADP ratios in MEPs and GMPs. (E) Protein expression of NDUFA9 and MTCO2 in MEPs and GMPs. TOMM20 and α-Tubulin were detected as loading controls for mitochondria and whole cells, respectively. (F, G) Mitochondrial ROS (F) and intracellular ROS (G) in MEPs, GMPs, and WBM cells. Values are presented as the means ± SD. Two-sided Welch’s *t*-test (C and D) was conducted to evaluate statistical significance. *P < 0.05; **P < 0.01.

One of the important functions of mitochondria is to synthesize iron-sulfur clusters and heme (Lane et al., 2015), both of which are molecular cargoes of iron for safe utilization of this redox-active element. Because erythroblasts need to synthesize a large amount of heme for hemoglobin production, we suspected that insufficient heme synthesis due to mitochondrial dysfunction might result in inhibition of erythroblast differentiation from *Cars2* IKO MEPs. However, neither heme content nor intracellular ferrous iron levels were changed in *Cars2* IKO cells (Figures S3A and S3B), suggesting that heme synthesis and iron availability are unlikely to link mitochondrial dysfunction to reduced erythroid commitment of *Cars2* IKO MEPs.

*Cars2* IKO GMPs showed similar changes to those of *Cars2* IKO MEPs except for MMP. Considering the limited effects of the *Cars2*^AINK^ mutation on myeloid cell differentiation *in vivo* and *in vitro* (see Figures 1D and 2A), GMPs are likely to be less susceptible to mitochondrial dysfunction than MEPs.

### Mitochondrial activity determines lineage choice in MEP differentiation

Based on the results of our genetic model of mitochondrial dysfunction, *Cars2* IKO mice, we pharmacologically disrupted mitochondrial function in wild-type MEPs. Addition of chloramphenicol (CAM), a specific inhibitor of mitochondrial translation, to the *in vitro* differentiation culture of MEPs markedly decreased and increased the frequency of erythroblasts and megakaryocytes, respectively (Figure 4A), suggesting that the production of ETC complex subunits encoded by mitochondrial DNA is required for erythroblast commitment. Inhibitors of each ETC complex (Figure S4A) gave similar results to those of CAM (Figure 4B and S4B). Mitochondrial inhibition by rotenone reproduced the results of *Cars2* IKO MEPs, showing a remarkable reduction in BFU-E formation without apparent changes in other colony types (Figure 4C). Intriguingly, FCCP, an uncoupler that depolarizes mitochondria, had no significant effect on MEP differentiation (Figures 4D, S4C and S4D). These results suggest that erythroid commitment of MEPs requires the electron transport via a functional ETC, but not MMP maintenance. Supporting this notion, complex V inhibition by oligomycin A, which reduces electron transport, inhibits erythroid commitment although MMP is maintained (Figure 4B and S4B). This confirms the necessity of electron transport for the erythroid commitment of MEPs, while demonstrating that MMP maintenance alone is insufficient.

**Figure 4.**
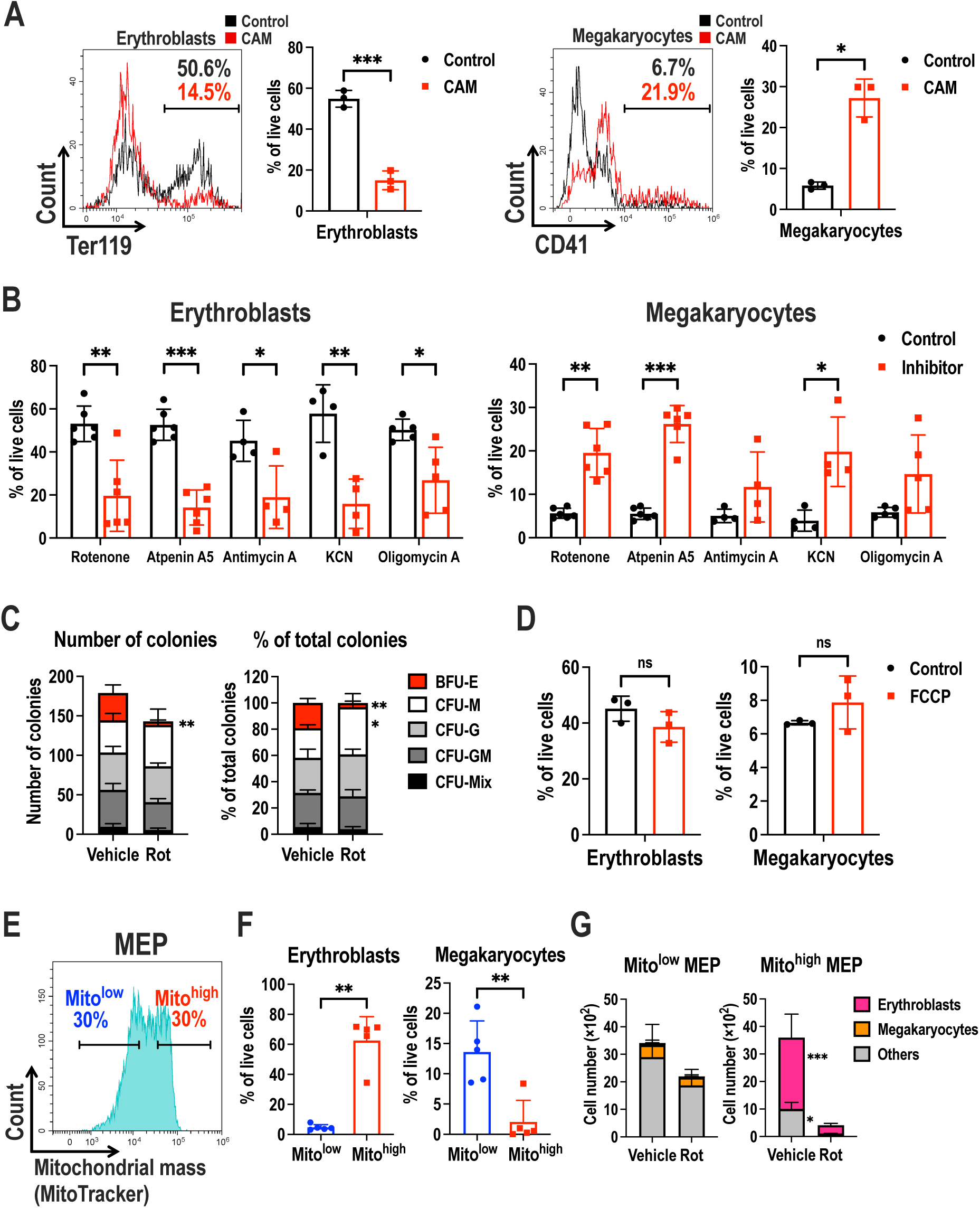
Mitochondrial activity determines lineage choice of MEP. (A) Representative histograms and ratios of erythroblasts and megakaryocytes in the MEP differentiation culture with or without chloramphenicol (CAM). (B) Ratios of erythroblasts and megakaryocytes in the MEP differentiation culture with or without mitochondrial ETC inhibitors shown in Figure S4A. (C) Colony counts (left) and ratios (right) of each colony type in colony-forming cell assay (*n* = 3) with or without rotenone (Rot). (D) Ratios of erythroblasts and megakaryocytes in the MEP differentiation culture with or without FCCP. (E) Representative histogram indicating MEP populations of bottom (Mito^low^) and top (Mito^high^) 30% of mitochondrial mass. (F) Ratios of erythroblasts and megakaryocytes differentiated from Mito^low^ and Mito^high^ MEPs. (G) Total live cell number of Mito^low^ and Mito^high^ MEPs cultured with or without rotenone (Rot) (*n* = 3). Values are presented as the means ± SD. Two-sided Welch’s *t*-test (A, B, D and F) and two-way ANOVA (C and G) were conducted to evaluate statistical significance. *P < 0.05; **P < 0.01; ***P < 0.001.

Finally, we asked whether an endogenous variation of mitochondrial activity that occurs within the MEP population plays a deterministic role in MEP lineage selection. We collected two subpopulations of MEPs, Mito^high^ MEPs as the top 30% and Mito^low^ MEPs as the bottom 30% of the MitoTracker signal corresponding to mitochondrial mass (Figure 4E). Surprisingly, the majority of Mito^high^ MEPs differentiated into erythroblasts, whereas the majority of Mito^low^ MEPs differentiated into megakaryocytes (Figures 4F and S4E). Mito^high^ MEPs and Mito^low^ MEPs were sensitive and insensitive, respectively, to mitochondrial ETC inhibition by rotenone (Figure 4G). This result suggests that Mito^high^ MEPs and Mito^low^ MEPs, which exhibit greater and lesser reliance on mitochondria, have distinct differentiation fates toward erythroblasts and megakaryocytes, respectively. Since rotenone treatment blocked BFU-E formation in the colony formation assay using whole MEPs, Mito^high^ MEPs are likely to represent cells at a primed stage toward erythroblasts to form BFU-E.

## Discussion

This study has demonstrated that mitochondria play an important role in the bipotential differentiation capacity of MEPs, and that MEPs exhibit heterogeneity in mitochondrial content. Specifically, higher mitochondrial content favors erythroid commitment, while lower content favors megakaryocyte commitment. For analyzing the *in vivo* contribution of mitochondria to cell fate decisions of HPCs, *Cars2* IKO mice proved to be an ideal model, as they exhibit suppressed mitochondrial function and altered MEP differentiation. The observed blockade of erythroid commitment in MEPs from *Cars2* IKO mice, which was reproduced by mitochondrial ETC inhibitors, prompted us to further investigate the differentiation potential of MEPs with varying mitochondrial content. These findings open new avenues for understanding the regulation of cell fate in HPCs, leading us to propose the concept of a “mitochondrial navigation” for cell fate decisions.

Interestingly, MEPs were particularly sensitive to the mitochondria dysfunction, whereas the granulo-monocyte and lymphoid lineages in *Cars2* IKO mice were only marginally affected. Previous studies demonstrated that mitochondria are essential for erythroblast maturation, primarily due to their role in heme synthesis and iron metabolism. Mutations in factors involved in these processes often lead to sideroblastic anemia (SA) (Ducamp & Fleming, 2019). However, in *Cars2* IKO BM cells, despite the mitochondrial dysfunction, the total heme and intracellular ferrous iron levels remained unchanged (see Figures S3A and S3B). This suggests that mitochondrial functions beyond heme and iron metabolism are necessary for MEPs to commit to erythroid differentiation. Consistently, heme synthesis and iron metabolism in MEPs are not as active as those in erythroblasts. An unanswered question is how mitochondria influence MEPs to select the erythroid lineage. Answering this could uncover a novel mechanism by which mitochondrially-derived signals initiate the gene expression cascade that drives erythroid-specific phenotype acquisition.

Recent studies have shown that the mitochondrially-anchored protein DELE1 conveys mitochondrial stress to the nucleus. In response to mitochondrial stress, DELE1 is released from mitochondria, binds to HRI and activates a transcription factor ATF4 (Quirós et al., 2017; Guo et al., 2020; Sekine et al., 2023). The DELE1-HRI-ATF4 pathway is recognized as a part of the integrated stress response (ISR) and is triggered by various forms of mitochondrial stress, such as inhibition of the ETC and the decoupling of MMP (Quirós et al., 2017). However, we believe that activation of the DELE1-HRI-ATF4 pathway is unlikely to explain the phenotypes observed in *Cars2* IKO MEPs. This is because MMP decoupling, which activates the DELE1-HRI-ATF4 pathway, does not inhibit the erythroid commitment of MEPs (see Figure 4D). This interpretation is further supported by the observation that MEPs derived from aged mice, which exhibit reduced MMP, can still produce both erythroblasts and megakaryocytes (unpublished observation). Although we cannot entirely exclude the possibility that ISR pathway contributes to the inhibition of erythroid commitment due to the mitochondrial dysregulation, it is likely that alternative mechanisms connecting mitochondria and the nucleus are involved.

The most compelling aspect of our discovery is that MEPs can be classified into two populations based on their mitochondrial content: Mito^high^ MEPs, which give rise to erythroid cells, and Mito^low^ MEPs, which generate megakaryocytes. In contrast, previous studies identified erythroid progenitors (ERPs) and megakaryocyte progenitors (MKPs) through mRNA expression profiles of transcription factors, as revealed by single-cell RNA-sequencing analyses (Paul et al., 2015; Lu et al., 2018). It has been suggested that high and low rates of cell cycling define ERPs and MKRs, respectively, based on the observation that inhibition of cell cycling promotes the priming of MEPs towards MKPs (Lu et al., 2018). If we assume that ERPs correspond to Mito^high^ MEPs and have a higher mitochondrial content, their elevated proliferation activity may be driven and supported mainly by the anabolic TCA cycle, which is coupled with mitochondrial ETC. This is consistent with the observation that erythroid commitment of MEPs requires electron transport in the mitochondrial ETC, rather than MMP maintenance for ATP production (see Figures 4B and 4D). Furthermore, mitochondria biogenesis is often enhanced in parallel with increased cell cycling and proliferation (Antico Arciuch et al., 2012).

In summary, our findings demonstrate that mitochondrial activity plays a regulatory role in the lineage choice of MEPs, highlighting a novel aspect of mitochondrial contribution to hematopoiesis. Although we have not yet examined pathological conditions involving increased erythroblast production, such as polycythemia vera, the insights gained from this study could enhance our understanding of mitochondria-related regulation in hematopoiesis and potentially inform new therapeutic strategies for polycythemia vera. Further research is needed to elucidate the molecular mechanisms by which mitochondria transmit signals to the nucleus, thereby guiding the binary fate decisions in MEPs.

## Materials and Methods

### Mice

All the mice utilized in this study were in the C57BL/6 genetic background. *Rosa*-CreER^T2^ mice and *Rosa*-LSL-tdTomato mice were purchased from Jackson laboratory. Establishment of *Cars2*^AINK^ mice was described previously (Matsunaga et al, 2023). To obtain *Cars2*^F^ allele, two loxP sequences were inserted to the up-stream and down-stream of exon 8, by using CRISPR-CAS9 system along with guide RNAs and a donor plasmid. The guide RNAs and the plasmid were injected into fertilized eggs derived from C57BL/6 strain mice. The obtained *Cars2*^F^ founder mice were backcrossed into C57BL/6 stain for three generations. Genotyping primer sets are shown in Table S1. To obtain *Cars2*^AINK/F^:*Rosa*-CreER^T2^:*Rosa*-LSL-tdTomato (*Cars2*IKO) and *Cars2*^+/F^:*Rosa*-CreER^T2^:*Rosa*-LSL-tdTomato (*Cars2*IC) mice, *Cars2*^F/F^:*Rosa-*CreER^T2^/*Rosa-*CreER^T2^ mice were crossed with *Cars2*^AINK/+^:*Rosa*-LSL-tdTomato/*Rosa*-LSL-tdTomato mice. To obtain *Cars2*^AINK/F^:*Rosa*-CreER^T2^ and *Cars2*^+/F^:*Rosa*-CreER^T2^ mice, *Cars2*^AINK/+^ mice were crossed with *Cars2*^F/F^:*Rosa-*CreER^T2^/*Rosa*-CreER^T2^ mice. Six to ten-week-old littermate mice were used for analysis following daily intraperitoneal injections of tamoxifen (75 mg/kg/day) (Sigma, Cat. No. T5648) for either 3 or 7 days. The mice were bred and housed under specific pathogen-free conditions with standard animal maintenance according to the regulations of *The Standards for Human Care and Use of Laboratory Animals of Tohoku University* (Tohoku University. 2007. Standards for human care and use of laboratory animals of Tohoku University. Tohoku University) and *The Guidelines for Proper Conduct of Animal Experiments* by the Ministry of Education, Culture, Sports, Science, and Technology of Japan (Science Council of Japan. 2006. Guidelines for proper conduct of animal experiments. Science Council of Japan, Ministry of Education, Culture, Sports, Science, and Technology of Japan).

### Peripheral blood analysis

PB was collected from the buccal region into 0.5 M EDTA and analyzed using a hemocytometer (Nihon Kohden Co.). For flow cytometric analysis, PB was treated with a hemolytic reagent for 30 min at room temperature and washed with phosphate-buffered saline (PBS).

### Flow cytometric analysis

BM cells were collected from femora, tibiae, ilia, and humeri. To analyze lineage-positive cells and HSPCs, the collected BM cells were treated with a hemolytic reagent for 20 min at room temperature. PB cells were prepared as described previously. For erythroid cell and megakaryocyte staining, the BM cells were stained with antibodies against Ter119 (TER-119), CD71 (RI7217), CD44 (IM7) and CD41 (MWReg30). For staining other lineage-positive cells, the hemolyzed BM cells were stained with antibodies against CD11b (M1/70), Gr-1 (RB6-8C5), B220 (RA3-6B2), and CD3e (145-2C11). For HSPC staining, lineage-positive cells were first labeled with biotin-conjugated antibodies against Ter119, CD11b, Gr-1, B220, CD8a, and CD4, followed by reaction with streptavidin-conjugated antibodies. After the removal of lineage-positive cells, the remaining lineage-negative cells were stained with antibodies against c-Kit (2B8), Sca-1 (D7), FcγR (93) and CD34 (RAM34). Dead cells were labeled with propidium iodide (PI) prior to analysis. Flow cytometric analysis was performed using CytoFLEX LX (Beckman coulter).

For cell sorting, hemolyzed lineage-positive and Sca-1-positive cells were labeled with biotin-conjugated antibodies and removed using MojoSort streptavidin nanobeads (BioLegend) according to the manufacturer’s instruction. The remaining lineage-negative cells were stained with antibodies against c-Kit, FcγR and CD34 for 30 min on ice. Before cell sorting, dead cells were labeled with PI or Zombie Yellow (BioLegend) following the manufacturer’s instructions. Flow cytometric cell sorting was performed using AriaⅡ (BD Bioscience) or CytoFLEX SRT (Beckman coulter).

### Bone marrow transplantation (BMT) assay

For non-competitive BMT, *Cars2*^+/F^:CreER^T2^:*Rosa*-LSL-tdTomato mice or *Cars2*^AINK/F^:CreER^T2^:*Rosa*-LSL-tdTomato mice were used as donors. BM cells were collected from femora, tibiae, ilia, and humeri of donor mice using PBS containing 1% penicillin/streptomycin (P/S) (Nacalai tesque, Kyoto, Japan). The BM cells (1 × 10^6^) were transplanted into lethally irradiated (X-ray, 7Gy) C57BL/6J mice via intravenous injection. Seven to eight weeks from post-BMT, tamoxifen (75 mg/kg/day) was administered via intraperitoneal injection for 3 days, and the recipient mice were analyzed on day 7 after the initiation of tamoxifen injection.

### MEP transplantation assay

For MEP transplantation, mice were sacrificed on day 7 after the start of tamoxifen injection. Sorted MEP cells (1 × 10^5^) were transplanted into non-irradiated wild-type mice (C57BL/6J) via intravenous injection. Eleven days post-transplantation, the recipient mice were analyzed, and tdTomato-positive cells were identified as donor-derived.

### Detection of mitochondrial mass, mitochondrial and intracellular ROS, and mitochondrial membrane potential

Mitochondrial mass, mitochondrial ROS, intracellular ROS, and MMP were measured by flow cytometry. Equal numbers of BM cells were stained with cell surface markers and washed with PBS. The cells were then incubated with Mitotracker Green (10 nM) (Invitrogen, Cat. No. M7514) for mitochondrial mass, MitoSOX Red (5 μM) (Invitrogen, Cat. No. M36008) for mitochondrial ROS, CellROX (250 nM) (Invitrogen, Cat. No. C10444) for intracellular ROS, and JC10 dye (AAT Bioquest) for MMP, following the manufacturer’s instructions. Briefly, the cells were incubated with each dye in PBS containing 50 μM verapamil for 30 min at 37℃, and washed once with PBS. Prior to analysis, dead cells were labeled with PI. The cells were analyzed using a CytoFLEX LX flow cytometer. For the MMP negative control, 50 nM FCCP (abcam) was added to the cells with JC10 dye, and the mixture was incubated for 30 min at 37℃.

### ATP/ADP ratio assay

Intracellular ATP/ADP ratio was measured using a luminescence detection using ADP/ATP ratio assay kit (Sigma, Cat. No. MAK135), following the manufacturer’s instructions. Briefly, MEPs or GMPs (5 × 10^4^) were sorted and plated on a 96-well plate with black walls. Reagents were added into the cells, and luminescence was detected using a Nifinie 200 PRO (TECAN).

### BrdU incorporation assay

For *in vivo* BrdU incorporation assay, BrdU (100 mg/kg) (Sigma, Cat. No. B9285-1G) was administered to mice via intraperitoneally injection. BM cells were isolated 24 hours after BrdU injection. Lineage-negative cells were prepared from the BM cells as described above and stained with antibodies against c-Kit, Sca-1, FcγR and CD34. Following staining, the incorporated BrdU was detected using BrdU assay kit (BD, Cat. No. 552598), according to the manufacturer’s instruction and analyzed using CytoFLEX LX flow cytometer.

### Colony forming cell assay

LK cells were seeded into 1 ml methylcellulose-based medium (Stemcell, MethoCult, Cat. No. M3434). After 7 days, the number of colonies was counted. For rotenone treatment, LK cells were seeded in 1 mL methylcellulose-based medium containing 50 nM rotenone.

### MEP culture assay

Sorted MEPs (1 × 10^4^ cells/ml) were cultured according to a previously reported protocol with some modifications (Gotoh et al., 2020). The MEPs were maintained in DMEM supplemented with 30% FBS (Nichirei Bioscicences Inx.), 1% P/S (Nacalai Tesque), insulin-transferrin-selenium-ethanolamine (ITS-X, Thermo Fisher), 2 U/ml erythropoietin (BioLegend), 25 ng/ml stem cell factor (BioLegend), 25 ng/ml IL-3 (BioLegend), 25 ng/ml IL-6 (BioLegend), 25 ng/ml thrombopoietin (BioLegend) and 0.1 mM 1-thioglycerol (Sigma) for 2 days. ETC inhibitors were added to the MEP culture medium from the beginning of the experiment. The inhibitors used were 10nM rotenone (Sigma, Cat. No. R8875), 1 μM atpenine A5 (Santa Cruz, Cat. No. SC-202475), 10 nM antimycin A (Sigma, Cat. No. A8674), 1mM potassium cyanide (KCN) (Wako, Cat. No. 166-03611), and 0.1 nM oligomycin A (Sigma, Cat. No. 75351). The uncoupler FCCP was added at a concentration of 50 nM to the MEP culture medium at the time of cell seeding and every 12 hours thereafter. The cells were harvested, stained with antibodies against Ter119 and CD41, and analyzed using a CytoFLEX LX flow cytometer.

### Immunoblot analysis

MEPs or GMPs were sorted from three mice (1.5-2.0 × 10^5^ cells/mouse) and pooled for immunoblotting. The cells were lysed in a buffer containing 200 mM Tris-HCl, 4% SDS, 20% glycerol, 9.3% DTT, and 0.001% bromophenol blue, sonicated and boiled at 95℃ for 10 minutes. Proteins were separated by SDS-PAGE and transferred onto PVDF membrane (Sigma, Immobilon-P, Cat. No. IPVH00010). The membrane was blocked with 5% skim milk in Tris-buffered saline containing 0.1% Tween 20 (TBS-T) and incubated overnight at 4℃ with primary antibodies against NDUFA9 (Abcam, Cat. No. ab14713), MTCO2 (ThermoFisher, Cat. No. PA5-88146), α-Tubulin (Sigma, Cat. No. T9026) and TOMM20 (Abcam, Cat. No. ab56783) in Signal Enhancer (Nacalai, Signal Enhancer HIKARI, Cat. No. 02272-74). After washing with TBS-T, the membranes were incubated with horseradish peroxidase (HRP)-conjugated anti-mouse or anti-rabbit IgG antibodies in TBS-T for 1 hour at room temperature and washed again with TBS-T. Finally, the membranes were treated with Chemi-Lumi One L (Nacalai Tesque) or ECL prime (GE Healthcare), and the signal was detected by Amercham ImageQuant 800 (Cytiva).

### Statistical analysis

All experiments were performed at least twice independently. Quantitative data are presented as the means ± SD and were analyzed using Welch’s t test. Two-way ANOVA was employed for chronological analyses of body weights and blood parameters, and analysis of colony forming cell. P values of <0.05 were considered statistically significant. All statistical analyses were performed using GraphPad Prism 9.

## Acknowledgements

We thank the Laboratory Animal Resource Center at University of Tsukuba for generating *Cars2* floxed mice, the Department of Medical Biochemistry for discussions, and the Biomedical Research Core of the Institute of Development, Aging and Cancer for technical supports. This work was supported by JSPS (grant numbers 24K11531 (M.S.), 21H05263 (T.A.), 24H00063 (T.A.), 21H04799 (H.M.), 21H05258 (T.A., H.M.), 21H05264 (H.M.) and 24H00605 (H.M.)) and JST SPRING, Grant Number JPMJSP2114 (E.S.). The funders had no role in the study design, data collection and analysis, decision to publish or manuscript preparation.

## Author contributions

E.S. conducted the experiments, analyzed the data and wrote early draft of the paper. S.M. designed the study, conducted the experiments, analyzed the data and wrote early draft of the paper. H.T. designed the study and interpreted the data. M.M., M.Y. and T.A. generated and provided critical materials and analyzed and interpreted the data. H.M. designed the study, conducted the experiments, supervised the research, analyzed the data and wrote early draft of the paper.

## Reference

Akaike T, Ida T, Wei FY, Nishida M, Kumagai Y, Alam MM, Ihara H, Sawa T, Matsunaga T, Kasamatsu S, Nishimura A, Morita M, Tomizawa K, Nishimura A, Watanabe S, Inaba K, Shima H, Tanuma N, Jung M, Fujii S, Watanabe Y, Ohmuraya M, Nagy P, Feelisch M, Fukuto JM, Motohashi H. Cysteinyl-tRNA synthetase governs cysteine polysulfidation and mitochondrial bioenergetics. Nat Commun. 2017 Oct 27;8(1):1177. doi: 10.1038/s41467-017-01311-y. PMID: 29079736; PMCID: PMC5660078.

Akaike T, Morita M, Ogata S, Yoshitake J, Jung M, Sekine H, Motohashi H, Barayeu U, Matsunaga T. New aspects of redox signaling mediated by supersulfides in health and disease. Free Radic Biol Med. 2024 Sep;222:539–551. doi: 10.1016/j.freeradbiomed.2024.07.007. Epub 2024 Jul 9. PMID: 38992395.

Akashi K, Traver D, Miyamoto T, Weissman IL. A clonogenic common myeloid progenitor that gives rise to all myeloid lineages. Nature. 2000 Mar 9;404(6774):193-7. doi: 10.1038/35004599. PMID: 10724173.

Alam MM, Kishino A, Sung E, Sekine H, Abe T, Murakami S, Akaike T, Motohashi H. Contribution of NRF2 to sulfur metabolism and mitochondrial activity. Redox Biol. 2023 Apr;60:102624. doi: 10.1016/j.redox.2023.102624. Epub 2023 Feb 2. PMID: 36758466; PMCID: PMC9941419.

Ansó E, Weinberg SE, Diebold LP, Thompson BJ, Malinge S, Schumacker PT, Liu X, Zhang Y, Shao Z, Steadman M, Marsh KM, Xu J, Crispino JD, Chandel NS. The mitochondrial respiratory chain is essential for haematopoietic stem cell function. Nat Cell Biol. 2017 Jun;19(6):614–625. doi: 10.1038/ncb3529. Epub 2017 May 15. PMID: 28504706; PMCID: PMC5474760.

Antico Arciuch VG, Elguero ME, Poderoso JJ, Carreras MC. Mitochondrial regulation of cell cycle and proliferation. Antioxid Redox Signal. 2012 May 15;16(10):1150–80. doi: 10.1089/ars.2011.4085. Epub 2012 Jan 13. PMID: 21967640; PMCID: PMC3315176.

Arinobu Y, Mizuno S, Chong Y, Shigematsu H, Iino T, Iwasaki H, Graf T, Mayfield R, Chan S, Kastner P, Akashi K. Reciprocal activation of GATA-1 and PU.1 marks initial specification of hematopoietic stem cells into myeloerythroid and myelolymphoid lineages. Cell Stem Cell. 2007 Oct 11;1(4):416–27. doi: 10.1016/j.stem.2007.07.004. PMID: 18371378.

Barayeu U, Sawa T, Nishida M, Wei FY, Motohashi H, Akaike T. Supersulfide biology and translational medicine for disease control. Br J Pharmacol. 2023 Oct 23. doi: 10.1111/bph.16271. Epub ahead of print. PMID: 37872133.

Chandel NS. Evolution of Mitochondria as Signaling Organelles. Cell Metab. 2015 Aug 4;22(2):204–6. doi: 10.1016/j.cmet.2015.05.013. Epub 2015 Jun 11. PMID: 26073494.

de Almeida MJ, Luchsinger LL, Corrigan DJ, Williams LJ, Snoeck HW. Dye-Independent Methods Reveal Elevated Mitochondrial Mass in Hematopoietic Stem Cells. Cell Stem Cell. 2017 Dec 7;21(6):725–729.e4. doi: 10.1016/j.stem.2017.11.002. Epub 2017 Nov 30. PMID: 29198942; PMCID: PMC5728653.

Doré LC, Crispino JD. Transcription factor networks in erythroid cell and megakaryocyte development. Blood. 2011 Jul 14;118(2):231–9. doi: 10.1182/blood-2011-04-285981. Epub 2011 May 26. PMID: 21622645; PMCID: PMC3138678.

Ducamp S, Fleming MD. The molecular genetics of sideroblastic anemia. Blood. 2019 Jan 3;133(1):59–69. doi: 10.1182/blood-2018-08-815951. Epub 2018 Nov 6. PMID: 30401706; PMCID: PMC6318428.

Fontenay M, Cathelin S, Amiot M, Gyan E, Solary E. Mitochondria in hematopoiesis and hematological diseases. Oncogene. 2006 Aug 7;25(34):4757–67. doi: 10.1038/sj.onc.1209606. PMID: 16892088.

Gong Y, Lan XP, Guo S. *IARS2*-related disease manifesting as sideroblastic anemia and hypoparathyroidism: A case report. Front Pediatr. 2023 Jan 10;10:1080664. doi: 10.3389/fped.2022.1080664. PMID: 36704128; PMCID: PMC9871752.

Gotoh K, Kunisaki Y, Mizuguchi S, Setoyama D, Hosokawa K, Yao H, Nakashima Y, Yagi M, Uchiumi T, Semba Y, Nogami J, Akashi K, Arai F, Kang D. Mitochondrial Protein Synthesis Is Essential for Terminal Differentiation of CD45^-^ TER119^-^Erythroid and Lymphoid Progenitors. iScience. 2020 Oct 7;23(11):101654. doi: 10.1016/j.isci.2020.101654. PMID: 33103089; PMCID: PMC7578749.

Guo X, Aviles G, Liu Y, Tian R, Unger BA, Lin YT, Wiita AP, Xu K, Correia MA, Kampmann M. Mitochondrial stress is relayed to the cytosol by an OMA1-DELE1-HRI pathway. Nature. 2020 Mar;579(7799):427-432. doi: 10.1038/s41586-020-2078-2. Epub 2020 Mar 4. PMID: 32132707; PMCID: PMC7147832.

Hoppe PS, Schwarzfischer M, Loeffler D, Kokkaliaris KD, Hilsenbeck O, Moritz N, Endele M, Filipczyk A, Gambardella A, Ahmed N, Etzrodt M, Coutu DL, Rieger MA, Marr C, Strasser MK, Schauberger B, Burtscher I, Ermakova O, Bürger A, Lickert H, Nerlov C, Theis FJ, Schroeder T. Early myeloid lineage choice is not initiated by random PU.1 to GATA1 protein ratios. Nature. 2016 Jul 14;535(7611):299-302. doi: 10.1038/nature18320. PMID: 27411635.

Ida T, Sawa T, Ihara H, Tsuchiya Y, Watanabe Y, Kumagai Y, Suematsu M, Motohashi H, Fujii S, Matsunaga T, Yamamoto M, Ono K, Devarie-Baez NO, Xian M, Fukuto JM, Akaike T. Reactive cysteine persulfides and S-polythiolation regulate oxidative stress and redox signaling. Proc Natl Acad Sci U S A. 2014 May 27;111(21):7606–11. doi: 10.1073/pnas.1321232111. Epub 2014 Apr 14. PMID: 24733942; PMCID: PMC4040604.

Ito K, Carracedo A, Weiss D, Arai F, Ala U, Avigan DE, Schafer ZT, Evans RM, Suda T, Lee CH, Pandolfi PP. A PML–PPAR-δ pathway for fatty acid oxidation regulates hematopoietic stem cell maintenance. Nat Med. 2012 Sep;18(9):1350–8. doi: 10.1038/nm.2882. PMID: 22902876; PMCID: PMC3566224.

Johnson KD, Soukup AA, Bresnick EH. GATA2 deficiency elevates interferon regulatory factor-8 to subvert a progenitor cell differentiation program. Blood Adv. 2022 Mar 8;6(5):1464–1473. doi: 10.1182/bloodadvances.2021006182. PMID: 35008108; PMCID: PMC8905696.

Kato H, Itoh-Nakadai A, Matsumoto M, Ishii Y, Watanabe-Matsui M, Ikeda M, Ebina-Shibuya R, Sato Y, Kobayashi M, Nishizawa H, Suzuki K, Muto A, Fujiwara T, Nannya Y, Malcovati L, Cazzola M, Ogawa S, Harigae H, Igarashi K. Infection perturbs Bach2- and Bach1-dependent erythroid lineage ‘choice’ to cause anemia. Nat Immunol. 2018 Oct;19(10):1059–1070. doi: 10.1038/s41590-018-0202-3. Epub 2018 Sep 24. PMID: 30250186.

Lane DJ, Merlot AM, Huang ML, Bae DH, Jansson PJ, Sahni S, Kalinowski DS, Richardson DR. Cellular iron uptake, trafficking and metabolism: Key molecules and mechanisms and their roles in disease. Biochim Biophys Acta. 2015 May;1853(5):1130–44. doi: 10.1016/j.bbamcr.2015.01.021. Epub 2015 Feb 4. PMID: 25661197.

Laslo P, Spooner CJ, Warmflash A, Lancki DW, Lee HJ, Sciammas R, Gantner BN, Dinner AR, Singh H. Multilineage transcriptional priming and determination of alternate hematopoietic cell fates. Cell. 2006 Aug 25;126(4):755–66. doi: 10.1016/j.cell.2006.06.052. PMID: 16923394.

Liu X, Zhang Y, Ni M, Cao H, Signer RAJ, Li D, Li M, Gu Z, Hu Z, Dickerson KE, Weinberg SE, Chandel NS, DeBerardinis RJ, Zhou F, Shao Z, Xu J. Regulation of mitochondrial biogenesis in erythropoiesis by mTORC1-mediated protein translation. Nat Cell Biol. 2017 Jun;19(6):626–638. doi: 10.1038/ncb3527. Epub 2017 May 15. PMID: 28504707; PMCID: PMC5771482.

Lu YC, Sanada C, Xavier-Ferrucio J, Wang L, Zhang PX, Grimes HL, Venkatasubramanian M, Chetal K, Aronow B, Salomonis N, Krause DS. The Molecular Signature of Megakaryocyte-Erythroid Progenitors Reveals a Role for the Cell Cycle in Fate Specification. Cell Rep. 2018 Nov 20;25(8):2083–2093.e4. doi: 10.1016/j.celrep.2018.10.084. Erratum in: Cell Rep. 2018 Dec 11;25(11):3229. doi: 10.1016/j.celrep.2018.11.075. PMID: 30463007; PMCID: PMC6336197.

Matsunaga T, Sano H, Takita K, Morita M, Yamanaka S, Ichikawa T, Numakura T, Ida T, Jung M, Ogata S, Yoon S, Fujino N, Kyogoku Y, Sasaki Y, Koarai A, Tamada T, Toyama A, Nakabayashi T, Kageyama L, Kyuwa S, Inaba K, Watanabe S, Nagy P, Sawa T, Oshiumi H, Ichinose M, Yamada M, Sugiura H, Wei FY, Motohashi H, Akaike T. Supersulphides provide airway protection in viral and chronic lung diseases. Nat Commun. 2023 Jul 25;14(1):4476. doi: 10.1038/s41467-023-40182-4. PMID: 37491435; PMCID: PMC10368687.

Murakami S, Shimizu R, Romeo PH, Yamamoto M, Motohashi H. Keap1-Nrf2 system regulates cell fate determination of hematopoietic stem cells. Genes Cells. 2014 Mar;19(3):239–53. doi: 10.1111/gtc.12126. Epub 2014 Jan 21. PMID: 24580727.

Murakami S, Kusano Y, Okazaki K, Akaike T, Motohashi H. NRF2 signalling in cytoprotection and metabolism. Br J Pharmacol. 2023 Sep 15. doi: 10.1111/bph.16246. Epub ahead of print. PMID: 37715470.

Paul F, Arkin Y, Giladi A, Jaitin DA, Kenigsberg E, Keren-Shaul H, Winter D, Lara-Astiaso D, Gury M, Weiner A, David E, Cohen N, Lauridsen FK, Haas S, Schlitzer A, Mildner A, Ginhoux F, Jung S, Trumpp A, Porse BT, Tanay A, Amit I. Transcriptional Heterogeneity and Lineage Commitment in Myeloid Progenitors. Cell. 2015 Dec 17;163(7):1663–77. doi: 10.1016/j.cell.2015.11.013. Epub 2015 Nov 25. Erratum in: Cell. 2016 Jan 14;164(1-2):325. PMID: 26627738.

Quirós PM, Prado MA, Zamboni N, D’Amico D, Williams RW, Finley D, Gygi SP, Auwerx J. Multi-omics analysis identifies ATF4 as a key regulator of the mitochondrial stress response in mammals. J Cell Biol. 2017 Jul 3;216(7):2027–2045. doi: 10.1083/jcb.201702058. Epub 2017 May 31. PMID: 28566324; PMCID: PMC5496626.

Riley LG, Cooper S, Hickey P, Rudinger-Thirion J, McKenzie M, Compton A, Lim SC, Thorburn D, Ryan MT, Giegé R, Bahlo M, Christodoulou J. Mutation of the mitochondrial tyrosyl-tRNA synthetase gene, YARS2, causes myopathy, lactic acidosis, and sideroblastic anemia--MLASA syndrome. Am J Hum Genet. 2010 Jul 9;87(1):52-9. doi: 10.1016/j.ajhg.2010.06.001. PMID: 20598274; PMCID: PMC2896778.

Riley LG, Rudinger-Thirion J, Schmitz-Abe K, Thorburn DR, Davis RL, Teo J, Arbuckle S, Cooper ST, Campagna DR, Frugier M, Markianos K, Sue CM, Fleming MD, Christodoulou J. LARS2 Variants Associated with Hydrops, Lactic Acidosis, Sideroblastic Anemia, and Multisystem Failure. JIMD Rep. 2016;28:49–57. doi: 10.1007/8904_2015_515. Epub 2015 Nov 5. PMID: 26537577; PMCID: PMC5059179.

Sekine H, Takeda H, Takeda N, Kishino A, Anzawa H, Isagawa T, Ohta N, Murakami S, Iwaki H, Kato N, Kimura S, Liu Z, Kato K, Katsuoka F, Yamamoto M, Miura F, Ito T, Takahashi M, Izumi Y, Fujita H, Yamagata H, Bamba T, Akaike T, Suzuki N, Kinoshita K, Motohashi H. PNPO-PLP axis senses prolonged hypoxia in macrophages by regulating lysosomal activity. Nat Metab. 2024 Jun;6(6):1108–1127. doi: 10.1038/s42255-024-01053-4. Epub 2024 May 31. PMID: 38822028.

Sekine Y, Houston R, Eckl EM, Fessler E, Narendra DP, Jae LT, Sekine S. A mitochondrial iron-responsive pathway regulated by DELE1. Mol Cell. 2023 Jun 15;83(12):2059–2076.e6. doi: 10.1016/j.molcel.2023.05.031. PMID: 37327776; PMCID: PMC10329284.

Sive JI, Göttgens B. Transcriptional network control of normal and leukaemic haematopoiesis. Exp Cell Res. 2014 Dec 10;329(2):255–64. doi: 10.1016/j.yexcr.2014.06.021. Epub 2014 Jul 8. PMID: 25014893; PMCID: PMC4261078.

Starck J, Cohet N, Gonnet C, Sarrazin S, Doubeikovskaia Z, Doubeikovski A, Verger A, Duterque-Coquillaud M, Morle F. Functional cross-antagonism between transcription factors FLI-1 and EKLF. Mol Cell Biol. 2003 Feb;23(4):1390–402. doi: 10.1128/MCB.23.4.1390-1402.2003. PMID: 12556498; PMCID: PMC141137.

Sun J, Ramos A, Chapman B, Johnnidis JB, Le L, Ho YJ, Klein A, Hofmann O, Camargo FD. Clonal dynamics of native haematopoiesis. Nature. 2014 Oct 16;514(7522):322-7. doi: 10.1038/nature13824. Epub 2014 Oct 5. PMID: 25296256; PMCID: PMC4408613.

Takeda H, Murakami S, Liu Z, Sawa T, Takahashi M, Izumi Y, Bamba T, Sato H, Akaike T, Sekine H, Motohashi H. Sulfur metabolic response in macrophage limits excessive inflammatory response by creating a negative feedback loop. Redox Biol. 2023 Sep;65:102834. doi: 10.1016/j.redox.2023.102834. Epub 2023 Jul 29. PMID: 37536084; PMCID: PMC10412850.

Yamamoto R, Morita Y, Ooehara J, Hamanaka S, Onodera M, Rudolph KL, Ema H, Nakauchi H. Clonal analysis unveils self-renewing lineage-restricted progenitors generated directly from hematopoietic stem cells. Cell. 2013 Aug 29;154(5):1112–1126. doi: 10.1016/j.cell.2013.08.007. PMID: 23993099.

Yu L, Takai J, Otsuki A, Katsuoka F, Suzuki M, Katayama S, Nezu M, Engel JD, Moriguchi T, Yamamoto M. Derepression of the DNA Methylation Machinery of the *Gata1* Gene Triggers the Differentiation Cue for Erythropoiesis. Mol Cell Biol. 2017 Mar 31;37(8):e00592–16. doi: 10.1128/MCB.00592-16. PMID: 28069743; PMCID: PMC5376630.

Yu WM, Liu X, Shen J, Jovanovic O, Pohl EE, Gerson SL, Finkel T, Broxmeyer HE, Qu CK. Metabolic regulation by the mitochondrial phosphatase PTPMT1 is required for hematopoietic stem cell differentiation. Cell Stem Cell. 2013 Jan 3;12(1):62–74. doi: 10.1016/j.stem.2012.11.022. PMID: 23290137; PMCID: PMC3632072.

Zhang T, Ono K, Tsutsuki H, Ihara H, Islam W, Akaike T, Sawa T. Enhanced Cellular Polysulfides Negatively Regulate TLR4 Signaling and Mitigate Lethal Endotoxin Shock. Cell Chem Biol. 2019 May 16;26(5):686–698.e4. doi: 10.1016/j.chembiol.2019.02.003. Epub 2019 Mar 7. PMID: 30853417.

