## Supplementary information for "Mitochondria Regulate the Cell Fate Decisions of Megakaryocyte-Erythroid Progenitors"

Eunkyu Sung[1], Shohei Murakami[1]*, Haruna Takeda[1], Masanobu Morita[2], Masayuki Yamamoto[3], Takaaki Akaike[2], Hozumi Motohashi[1,4]*.

[1]Department of Medical Biochemistry, Tohoku University Graduate School of Medicine; Sendai, Japan

[2]Department of Environmental Medicine and Molecular Toxicology, Tohoku University Graduate School of Medicine; Sendai, Japan.

[3]Department of Biochemistry and Molecular Biology, Tohoku Medical Megabank Organization, Tohoku University, Sendai, Japan.

[4]Department of Gene Expression Regulation, IDAC, Tohoku University; Sendai, Japan.

*Corresponding authors:
Shohei Murakami
Department of Medical Biochemistry, Tohoku University Graduate School of Medicine.
2-1 Seiryo-machi, Aoba-ku, Sendai 980-8575, Japan


Hozumi Motohashi
Department of Medical Biochemistry, Tohoku University Graduate School of Medicine.
2-1 Seiryo-machi, Aoba-ku, Sendai 980-8575, Japan


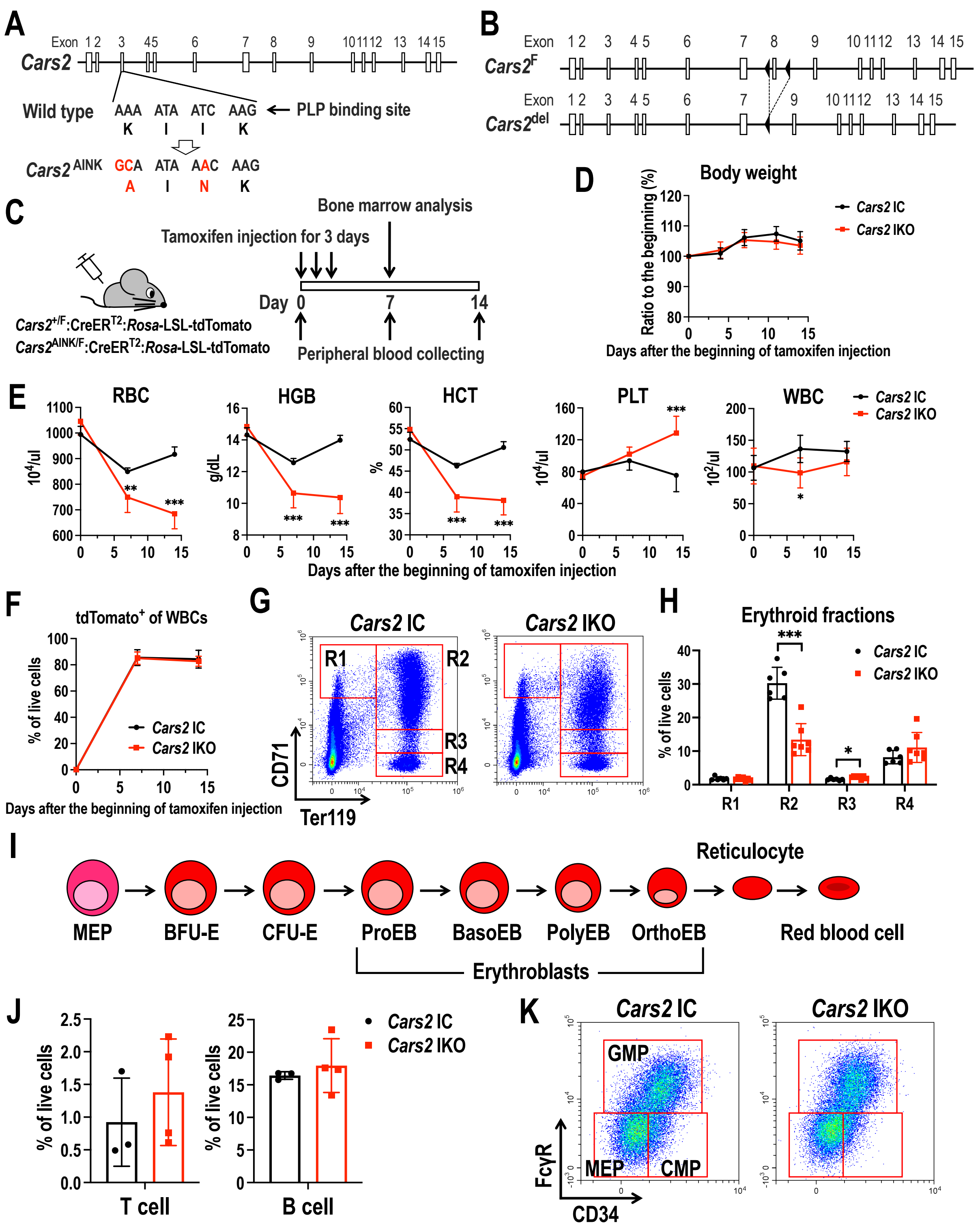


**Supplementary Figure S1. An anemia-like phenotype in *Cars2* IKO mice. Related to Figure 1.**

(A) *Cars2* gene locus and sequence of wild-type and *Cars2*$^{\mathrm{AINK}}$ mutant alleles at KIIK motif in exon 3, which is a binding site of pyridoxal 5'-phosphate (PLP).

(B) Floxed *Cars2* allele (*Cars2*$^{\mathrm{F}}$) and deleted *Cars2* allele (*Cars2*$^{\mathrm{del}}$). Black arrowheads indicate loxP sequences. Exon 8 is deleted by Cre recombinase.

(C) Experimental design of inducible gene manipulation by tamoxifen injection and analysis. Tamoxifen was intraperitoneally injected to *Cars2* IC (*Cars2*$^{+/F}$:*Rosa*-CreER$^{T2}$:*Rosa*-LSL-tdTomato) and *Cars2* IKO (*Cars2*$^{AINK/F}$:*Rosa*-CreER$^{T2}$:*Rosa*-LSL-tdTomato) mice for 3 days. Peripheral blood was collected from the mice at day 0, day 7, and day 14 from the beginning of tamoxifen injection. Bone marrows from the mice were analyzed at day 7. Cre-mediated recombination was monitored by tdTomato fluorescence.
(D) Body weights of *Cars2* IC and *Cars2* IKO mice for 14 days from the beginning of tamoxifen injection ($n$ = 6).
(E) Blood counts in *Cars2* IC and *Cars2* IKO mice (n = *6*). To study changes in the blood counts over time, blood was taken from the same mice 3 times.
(F) Ratios of tdTomato-positive cells in peripheral blood from *Cars2* IC and *Cars2* IKO mice (n = *6*), by which recombination efficiency was monitored.
(G) Representative plots of CD71 versus Ter119 in live BM cells.
(H) Ratios of erythroblast fractions (R1-R4 in panel G) in live BM cells.
(I) Differentiation stages of erythroid lineage.
(J) Ratios of T cells and B cells in BM.
(K) Representative plots of FcγR versus CD34 in Lin$^{-}$Sca1$^{-}$cKit$^{+}$ (LK) cells.
Values are presented as the means ± SD. Two-way ANOVA (D and E) and two-sided Welch's *t*-test (H and J) were conducted to evaluate statistical significance. *P < 0.05; **P < 0.01; ***P < 0.001.

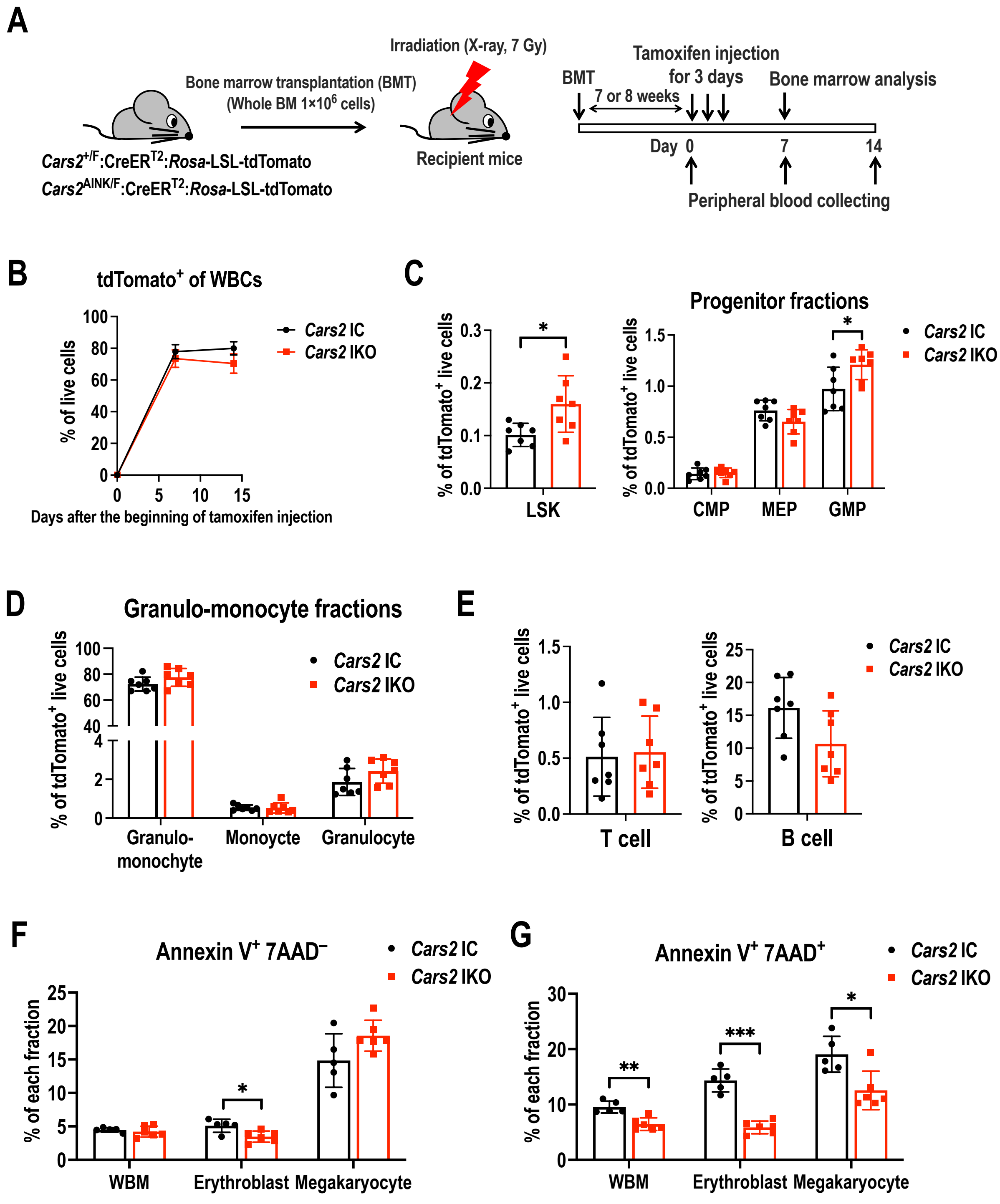


**Supplementary Figure 2. Non-competitive BMT. Related to Figure 1.**

(A) Experimental design for non-competitive BMT. Whole BM cells ($1.0 \times 10^6$) were taken from *Cars2* IC (*Cars2*$^{+/F}$:*Rosa*-CreER$^{T2}$:*Rosa*-LSL-tdTomato) and *Cars2* IKO (*Cars2*$^{AINK/F}$:*Rosa*-CreER$^{T2}$:*Rosa*-LSL-tdTomato) mice and transplanted into lethally-irradiated (X-ray, 7Gy) wild-type mice. Tamoxifen was injected into the recipient mice for 3 days from 7-8 weeks after the transplantation.

(B) Ratios of tdTomato-positive cells in peripheral blood from the recipient mice, by which

recombination efficiency and BM reconstitution efficiency were monitored.
(C-E) Ratios of donor-derived cells evaluated by the frequency of tdTomato-positive cells. LSK and progenitor fractions (C), myeloid cells (D) and lymphocytes (E).
(F-G) Percentages of Annexin $V^{+}7AAD^{-}$ (F) and Annexin $V^{+}7AAD^{+}$ cells (G) in WBM cells, erythroblasts (R2 fraction in Figure S1G) and megakaryocytes.
Values are presented as the means ± SD. Two-sided Welch's *t*-test (C-G) was conducted to evaluate statistical significance. *P < 0.05; **P < 0.01; ***P < 0.001.

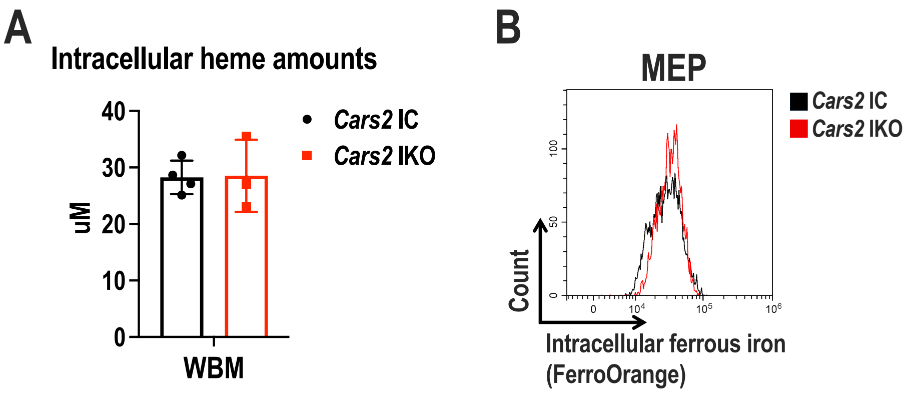


**Supplementary Figure 3. Intracellular heme contents and ferrous iron levels. Related to Figure 4.**

(A) Total intracellular heme amounts in whole BM cells.

(B) Representative histogram showing intracellular ferrous iron levels in MEPs.

Values are presented as the means ± SD. Two-sided Welch's *t*-test (A) was conducted to evaluate statistical significance.

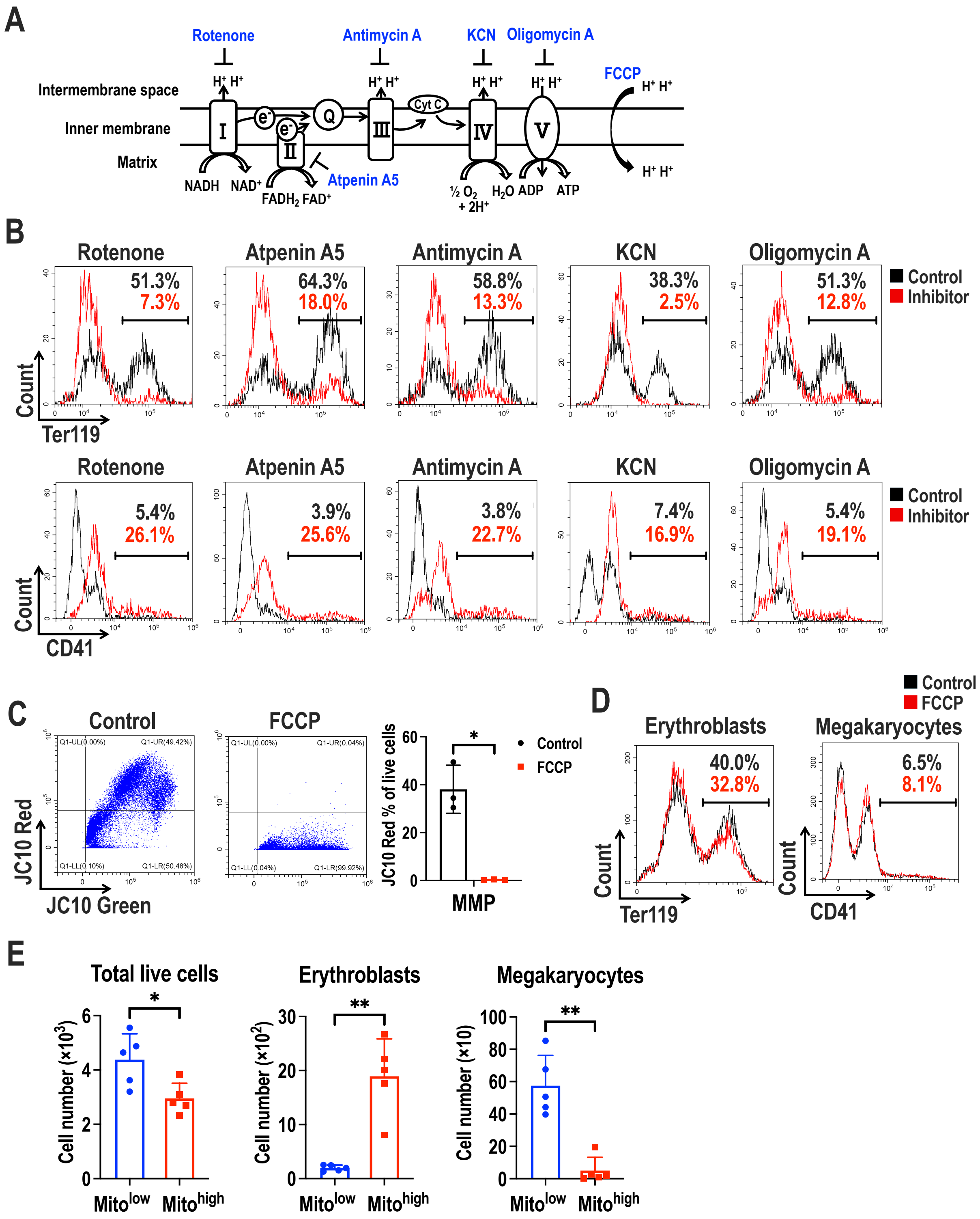


**Supplementary Figure 4. Effects of mitochondrial activities on lineage selection of MEPs. Related to Figure 4.**

(A) ETC complexes and their inhibitors.

(B) Representative histograms to detect erythroblasts (Ter119$^{+}$) and megakaryocytes (CD41$^{+}$) in the differentiation culture of MEPs with or without ETC inhibitors.

(C) Representative dot plots (left) and ratios of JC10-Red-positive cells (right) in the

differentiation culture of MEPs treated with FCCP.
(D) Representative histograms to detect erythroblasts ($Terr19^+$) and megakaryocytes ($CD41^+$) in the differentiation culture of MEPs with or without FCCP.
(E) Counts of total live cells, erythroblasts and megakaryocytes differentiated from $Mito^{low}$ and $Mito^{high}$ MEPs.
Values are presented as the means ± SD. Two-sided Welch's *t*-test (C and E) was conducted to evaluate statistical significance. *P < 0.05; **P < 0.01.

Supplementary Table S1. Genotyping primers.

| **Genes** | **Forward/Reverse** | **Primer sequence (5'→3')** |
|---|---|---|
| *Cars2*$^{F}$ | Forward | TTTAGGGCAGGGCCCTCGAC |
| | Reverse | CTAGCCAGGAGGCCGCATAA |
| *Cars2*$^{AINK}$ | Forward | ACCTTTAATCTCAGCACTTGGGAG |
| | Reverse | TTACCTCGTTAGCTCTCTTGTTTATTGC |
| *Rosa*-Cre | Forward | ACGTTCACCGGCATCAACGT |
| | Reverse | CTGCATTACCGGTCGATGCA |
| *Rosa*-tdTomato | Forward | GGCATTAAAGCAGCGTATCC |
| | Reverse | CTGTTCCTGTACGGCATGG |
